# Cyclic peptide structure prediction and design using AlphaFold

**DOI:** 10.1101/2023.02.25.529956

**Authors:** Stephen A. Rettie, Katelyn V. Campbell, Asim K. Bera, Alex Kang, Simon Kozlov, Joshmyn De La Cruz, Victor Adebomi, Guangfeng Zhou, Frank DiMaio, Sergey Ovchinnikov, Gaurav Bhardwaj

**Affiliations:** Molecular and Cell Biology program, University of Washington, Seattle, WA, USA; Institute for Protein Design, University of Washington, Seattle, WA, USA; Department of Biochemistry, University of Washington, Seattle, WA, USA; Department of Medicinal Chemistry, University of Washington, Seattle, WA, USA; John Harvard Distinguished Science Fellowship, Harvard University, Cambridge, MA, USA; FAS Division of Science, Harvard University, Cambridge, MA, USA

## Abstract

Deep learning networks offer considerable opportunities for accurate structure prediction and design of biomolecules. While cyclic peptides have gained significant traction as a therapeutic modality, developing deep learning methods for designing such peptides has been slow, mostly due to the small number of available structures for molecules in this size range. Here, we report approaches to modify the AlphaFold network for accurate structure prediction and design of cyclic peptides. Our results show this approach can accurately predict the structures of native cyclic peptides from a single sequence, with 36 out of 49 cases predicted with high confidence (pLDDT > 0.85) matching the native structure with root mean squared deviation (RMSD) less than 1.5 Å. Further extending our approach, we describe computational methods for designing sequences of peptide backbones generated by other backbone sampling methods and for *de novo* design of new macrocyclic peptides. We extensively sampled the structural diversity of cyclic peptides between 7–13 amino acids, and identified around 10,000 unique design candidates predicted to fold into the designed structures with high confidence. X-ray crystal structures for seven sequences with diverse sizes and structures designed by our approach match very closely with the design models (root mean squared deviation < 1.0 Å), highlighting the atomic level accuracy in our approach. The computational methods and scaffolds developed here provide the basis for custom-designing peptides for targeted therapeutic applications.

## INTRODUCTION

Deep learning (DL) methods, such as AlphaFold and RoseTTAFold, have demonstrated remarkable accuracy in predicting the three-dimensional structure of proteins from their amino acid sequences (*1, 2*) and have now been successfully applied to predict the protein structures and protein-protein interaction networks at the proteome scale (*3, 4*). These structure prediction networks have also spurred the development of DL methods for designing new proteins with diverse shapes, sizes, and functions (*5–11*). However, these computational design studies have been primarily limited to larger proteins composed of canonical amino acids. While benchmarking studies show limited applicability of these methods for predicting structures of small peptides and peptide-protein complexes (*12, 13*), new approaches are required to adapt these algorithms for cyclic peptides. Macrocyclization is common in biologically active natural products and therapeutic peptide discovery campaigns as it confers several structural, stability, and permeability advantages. The lack of free termini in cyclic peptides makes them more resistant to exoproteases and peptidases, and despite the lack of regular secondary structures, the cyclic constraints can lock small peptides into stable folds (*14, 15*). Macrocycles also offer new opportunities to disrupt intracellular protein-protein interactions that play key roles in many biological processes and are difficult to drug with small molecules and antibodies (*15–17*). We previously described methods for accurate computational design of peptide macrocycles using the kinematic closure (KIC) algorithm to sample cyclic peptide backbones, followed by Rosetta sequence design (*18, 19*). However, that approach is computationally expensive and requires massive sampling of cyclic backbones and sequence design to generate promising design models. Here, we sought to develop DL-based methods for rapid and accurate structure prediction and design of cyclic peptides.

The small number of high-resolution structures for macrocycles makes it difficult to train a new macrocycle-specific DL model from scratch. Alternatively, new DL models can be trained on design models from previously described Rosetta and molecular dynamics-based approaches (*18–22*). However, the accuracy and performance of such a model would be limited by the accuracy of the methods used to generate the training data. Alternatively, pre-trained networks like AlphaFold and RoseTTAFold can be modified to recognize macrocyclization and benchmarked to determine their accuracy at predicting cyclic protein and peptide structures. In our previous KIC-based approach, we noted that cyclic peptides are primarily composed of canonical motifs and turn types that are also common in the loop regions of larger proteins. Indeed, the Rosetta energy function, which is derived from crystal structures of large proteins, is able to correctly capture these motifs during peptide design (*12, 18*). Further, recent benchmarking studies have shown that AlphaFold is able to predict the structures of short peptide ligands in apo as well as bound states (*12, 13, 23*). Therefore, we reasoned that information encoded in the AlphaFold network would be adequate to accurately predict and design macrocycles if cyclic constraints and a positional encoding invariant to cyclic permutations of the sequence can be enforced appropriately. Here, we describe an approach to encode cyclization as an input positional encoding for AlphaFold and test the accuracy of these changes in predicting the structures for cyclic peptides available in the Protein Data Bank (PDB). Next, we report an approach to redesign sequences of macrocyclic backbones using AlphaFold to improve their propensity to fold into the designed structures. Finally, we describe an approach for successfully hallucinating new macrocycles from scratch and use it to enumerate the rich structural diversity of structured macrocycles between 7–13 amino acids.

### Structure Prediction of Cyclic Peptides

We set out to expand AlphaFold for the structure prediction of cyclic peptides by modifying the inputs for relative positional encoding. For a linear peptide, the relative positional encoding defines the sequence separation between amino acids, with adjacent residues having a sequence separation of one and the N- and C-termini separated by the length -1 of the peptide (Figure 1A). To apply cyclic constraints, we defined and applied a custom N x N cyclic offset matrix that introduces the circularization to the relative positional encoding and changes the sequence separation between terminal residues for a peptide of length N to be one, too (Figure 1B). The relative positional encoding, after one-hot encoding and linear projection, is added to the pairwise feature within the evoformer module of AlphaFold network. Without this encoding, attention layers are permutation and order invariant. We implemented these changes in the ColabDesign framework, which implements AlphaFold for structure prediction and design (*24*), and termed it AfCycDesign in this work. We initially tested this on randomly selected cyclic peptide sequences from Protein Data Bank (PDB), and found the outputs from these initial tests showed correct peptide bond connection and geometry of the terminal residues without introducing distortions in the rest of the peptide structure (Supplementary Figure S1A). We next tested whether the output predictions would change if the circular permutations of a sequence were given as the inputs. The output structures for all circularly permuted sequences were very similar to each other (Supplementary Figure S1B).

**Figure 1:**
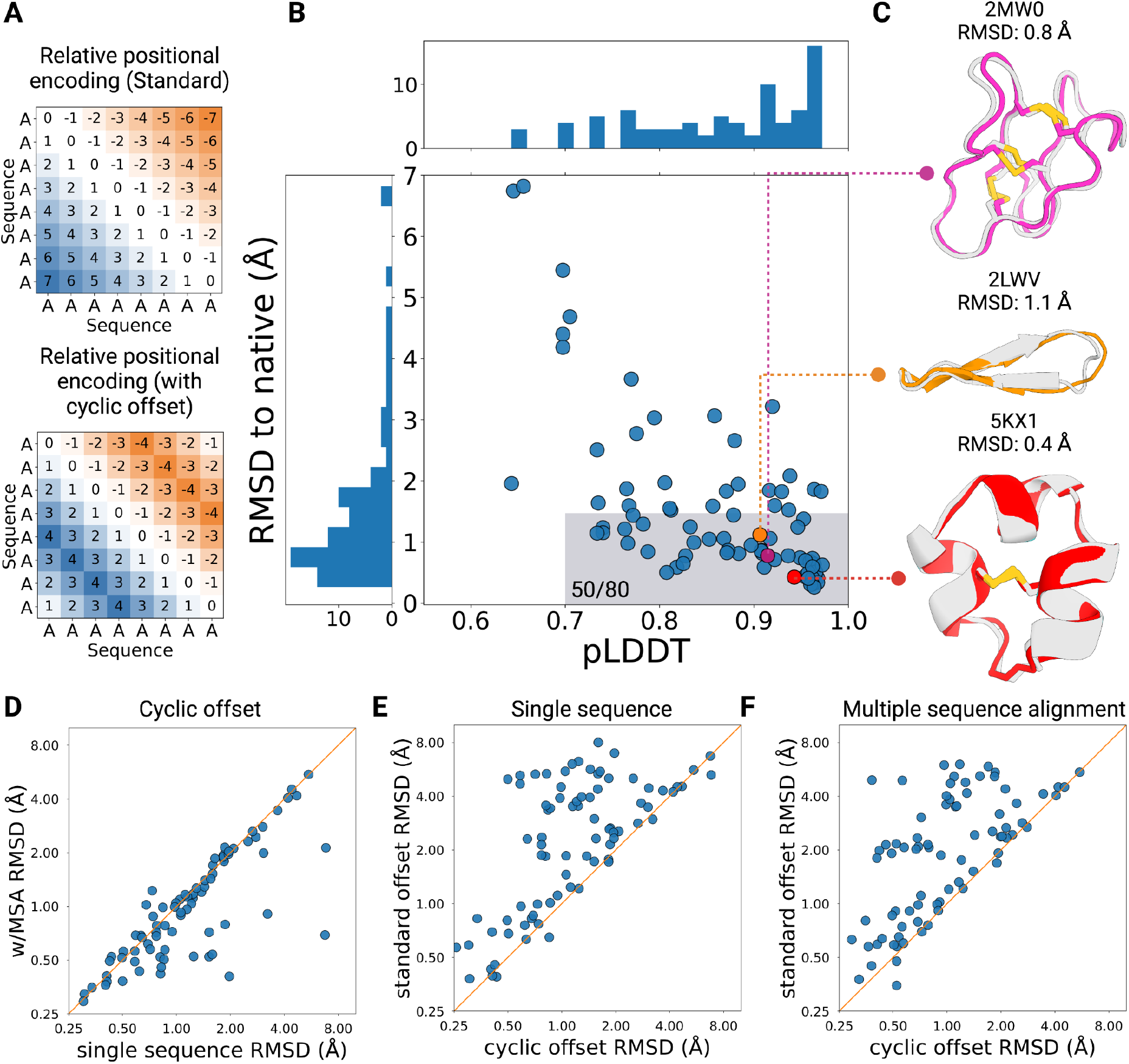
Structure prediction of native cyclic peptides using AfCycDesign. (A) Example of the relative positional encoding for a hypothetical eight residue peptide. Standard encoding in AfDesign shows sequence separation between residue positions for a linear peptide, with the termini being the maximum distance from each other. Application of the cyclic offset in AfCycDesign changes this behavior such that the termini are connected to each other. (B) AfCycDesign predictions of 80 cyclic peptides from the Protein Data Bank. The highlighted area covers good confidence and accurate predictions with pLDDT > 0.7 and RMSD < 1.5 Å. (C) Three representative predictions demonstrating the diverse topologies in structures predicted correctly (RMSD < 1.5 Å) with high confidence (pLDDT > 0.85) by AfCycDesign. The experimentally-determined structure is shown in gray and predicted structures from AfCycDesign are shown in magenta, orange, or red colors. (D) Comparison of the accuracy (RMSD to the native structure) of AfCycDesign prediction with cyclic offset from a single sequence and with MSA. (E) Comparisons of the accuracy of single sequence prediction with and without the cyclic offset. (F) Comparisons of the accuracy of MSA-based prediction with and without the cyclic offset.

Next, we assessed the accuracy of AfCycDesign at predicting the structures of diverse cyclic peptides deposited in the PDB. We collected 80 NMR structures from the PDB composed of canonical amino acids and sequence lengths of less than 40 amino acids. These were not in the training set of AlphaFold, since the training excluded NMR structures and short peptides with lengths less than 16 amino acids. These structures cover a broad range of topologies, with diverse sizes, secondary structures, sequences, and functions (Supplementary Table S1). Notably, many of the peptides in our test set, such as diverse plant-derived cyclotides or circular knottin folds, consist of multiple cysteine residues and disulfide bonds (*25*). The multiple possible disulfide-bond connectivities — 3 possible connectivities for four cysteines, and 15 for six cysteines — posed challenges for our previous methods, and disulfide connectivities had to be defined explicitly(*20*). We predicted the structure for each sequence in the test set using AfCycDesign and evaluated two metrics: the backbone heavy atom RMSD to the experimentally-determined structures and the predicted local distance difference test (pLDDT), a structure prediction confidence metric, from all five output models from AfCycDesign (see Methods for details). Overall, the predictions from AfCycDesign are close to the experimentally-determined structures with median pLDDT and RMSD of 0.88 and 1.13 Å, respectively (Figure 1B). In 50 out of 80 test cases, the predicted structures showed good confidence (pLDDT > 0.7) and backbone RMSD of less than 1.5 Å to the native structures. Notably, in 49 cases where AfCycDesign predicted structures with high confidence (plDDT > 0.85), 73% (n = 36) were predicted correctly with backbone heavy atom RMSD to native structure < 1.5 Å, indicating that pLDDT scores can be used to filter for accurate predictions for cyclic peptides. Notably, the correctly predicted structures were not limited to a specific class of peptides or topology and covered diverse sizes and topologies, including disulfide-rich cyclic peptides, small cyclic β-sheets, and peptides with very short α-helical motifs (Figure 1C). In 14 cases, the predicted structure was very close to the experimental structure (backbone RMSD < 1.5 Å); however, AfCycDesign had lower confidence in those predictions (pLDDT < 0.85) (Figure 1B). The lower confidence in these predictions stemmed from the loop regions that are also flexible in the NMR ensembles (Supplementary Figure S2). Despite there being no additional constraints placed on disulfide connectivity, correct bond connectivity was formed for most cases predicted with high confidence, which bodes well for structure prediction of knottins, conopeptides, cyclotides, and other classes of disulfide-rich peptides with many available sequences but very few experimentally determined structures. Adding multiple sequence alignments (MSA) during the predictions further improved the accuracy, with 58 out 80 cases correctly predicted (backbone heavy atom RMSD < 1.5 Å) with good confidence (pLDDT > 0.7), and the overall accuracy of prediction improved for a substantial number of structures (Figure 1D). In contrast, removing the cyclic offsets during single sequence or MSA-based predictions showed significantly decreased ability to predict the correct structures (Figure 1E-F). Taken together, these data highlight the promising accuracy of our AfCycDesign approach for predicting the structures of cyclic peptides with diverse sequences and structures.

### Sequence Redesign of Cyclic Peptides

We next applied cyclic relative positional encoding for designing amino acid sequences of cyclic peptides with AfCycDesign. We reasoned that such an approach would be useful in identifying new amino acid sequences that improve the folding propensity for a given backbone from natural-occurring peptides or generated using other backbone sampling approaches. To achieve this, we introduced cyclic offsets to the AfDesign approach previously implemented in ColabDesign. The goal of this approach is to find sequences predicted to fold into the desired backbone with high confidence by AlphaFold. This approach starts by predicting the distogram from a random sequence using the AlphaFold network and iteratively optimizing sequence at each subsequent step to minimize the difference between the predicted structure at that step and the desired backbone. The sequence optimization is guided by the difference (or the categorical cross-entropy) between the predicted distrogram (a tensor that contains a binned distribution of distances for every pair of residues) and one extracted from the desired structures (Supplementary Figure S3) (see Methods). This was shown to be a good proxy for maximizing the confidence of AlphaFold and minimizing the difference between the predicted and desired structure (*26*).

We set out to design peptide scaffolds suitable for targeting the helix-helix interactions common at protein-protein interfaces (*27*). We generated 457,615 backbones for 13-mer cyclic peptides that all included a short seven amino acid helix using our previously described Rosetta macrocycle design approach (*18*). To identify all the unique shapes in this large-scale run, we clustered the resulting backbones using a torsion-based binning approach: a bin string representing the structure was generated where each amino acid was assigned a bin based on the ɸ, ψ, and ω torsion angles; bins A and B refer to the α and β regions of the Ramachandran plot, while the bins X and Y refer to the mirrored regions in the positive ɸ region of the plot. Circular permutations of a bin string were also grouped into the same structural cluster. We identified 29,249 clusters with unique bin strings and selected one backbone, RRR13.1 (representing bin sequence: AAAAAAXBYBBAB), for redesign as the Rosetta-designed sequence for the same backbone had a small energy gap (ΔE < 2kcal/mol) in its energy landscapes (Figure 2A) (*18, 20*). The sequence designed by AfCycDesign differed significantly from the Rosetta-designed sequence, with 12 mutations in the sequence and only a singular alanine in the core of the peptide being conserved. We first evaluated the folding propensity of the AlphaFold-designed sequence, RAR13.1, *in silico* by calculating the structure-energy landscape using Rosetta cyclic peptide prediction methods (*18, 20*). The AlphaFold sequence converges to the designed structure as its lowest energy conformation with a larger energy gap (ΔE ∼ 6.0 kcal/mol) between the designed structure and alternative conformations (Figure 2A,B). To validate whether the AlphaFold-designed sequence actually folds into the designed structure, we determined its three-dimensional structure using racemic high-resolution X-ray crystallography and compared it to the computationally-designed model. The X-ray crystal structure was very close to the design model, with Cα RMSD of 0.2 Å and 10 out of the 13 sidechain rotamers in the X-ray crystal structure matching the designed model (Figure 2C).

**Figure 2:**
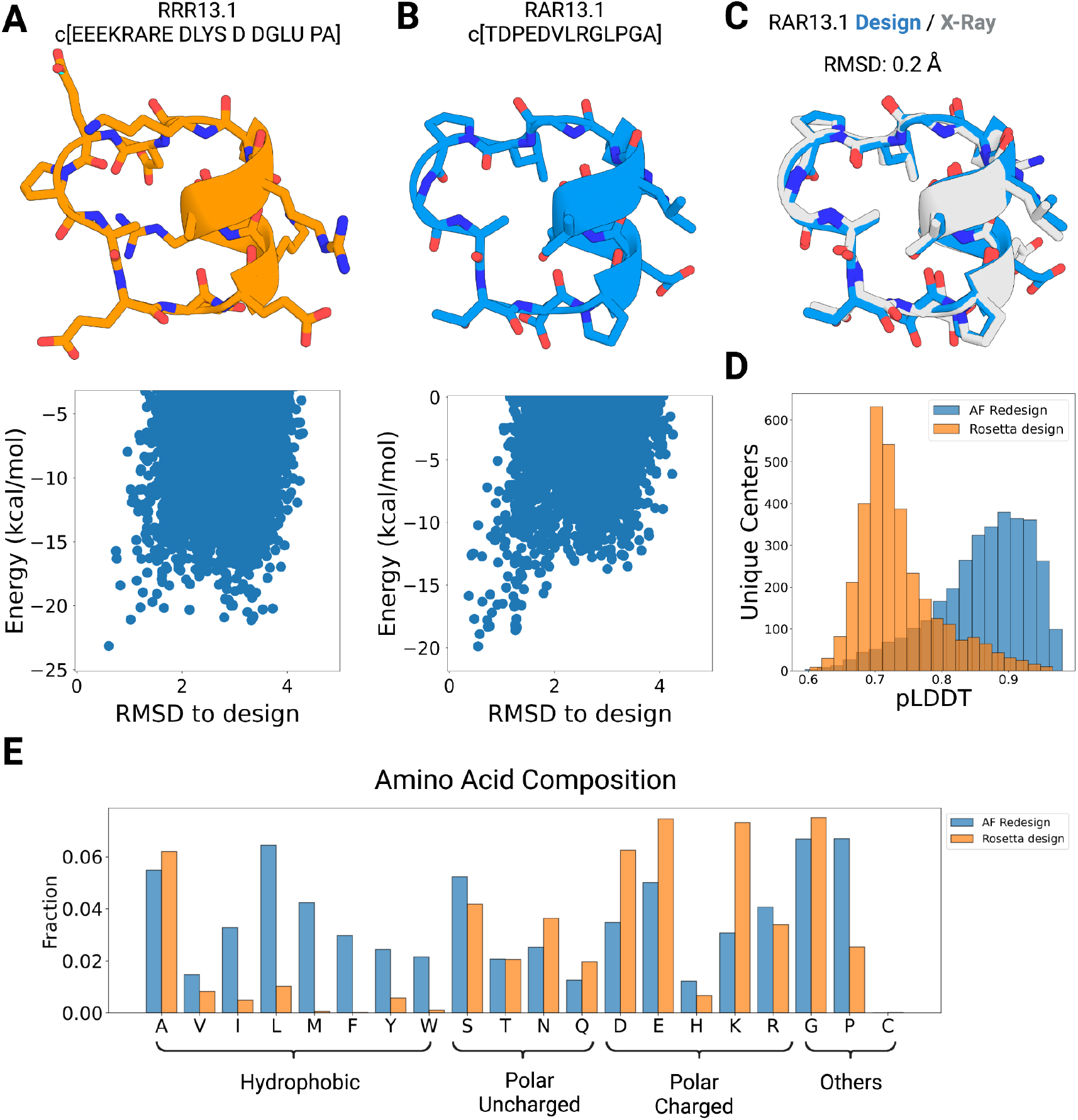
Sequence design of cyclic peptide backbones using AfCycDesign. (A) Sequence and design model for Rosetta-designed 13-residue cyclic peptide, RRR13.1. L-amino acids in the sequence are denoted by their one-letter codes, while D-amino acids are written with four-letter codes. The predicted energy landscape calculated by Rosetta cyclic peptide prediction methods is shown below the structure. (B) Sequence and design model for 13-residue cyclic peptide, RAR13.1, as designed by AfCycDesign. L-amino acids in the sequence are denoted by their one-letter codes. The predicted energy landscape calculated by Rosetta is shown below the structure. (C) Alignment of the RAR13.1 design model (blue) and the high-resolution X-ray crystal structure (gray) shows a very close match with Cα RMSD of 0.2 Å. (D) Distribution of pLDDT scores for sequences designed by Rosetta (orange) and AfCycDesign (blue). Each population was designed from the same backbones representing 3274 unique structural clusters of 13-residue peptides. (E) Frequency of each amino acid in sequences designed by Rosetta (orange) and AfCycDesign (blue) for backbones from 3274 unique structural clusters of 13-residue peptides.

Given the successful structure validation of AlphaFold-designed RAR13.1, we decided to redesign representative peptides from the 3,274 unique structural clusters, selected from our large-scale backbone sampling runs to have calculated Rosetta energy of less than 0 kcal/mol with poly alanine (or D-alanine) threaded on the backbones. In parallel, we also designed the selected backbones with Rosetta and compared them to sequences generated by AfCycDesign. As expected, the sequences designed by AfCycDesign had a better pLDDT score distribution than Rosetta-designed sequences for the same backbones (Figure 2D). While only 63 clusters had pLDDTs > 0.9 for Rosetta-designed sequences; 1145 clusters had pLDDTs > 0.9 among sequences from AfCycDesign (Figure 2D). However, we had to limit Rosetta design approach to the canonical 20 amino acids (as was required for pLDDT calculation), instead of allowing for heterochiral design as done in our previous work (*18–20*). In addition to comparing the structure prediction confidence metrics, we also explored the differences in amino acid composition and chemical properties for sequences designed by AfCycDesign and Rosetta. Sequences designed by AfCycDesign were generally more hydrophobic and included more prolines, compared to the Rosetta-designed sequences for the same backbones (Figure 2E). Overall, our *in silico* and experimental results demonstrate that AfCycDesign can be leveraged to design sequences for cyclic peptide backbones that fold into the desired structures. More broadly, the AfCycDesign approach is complementary to other methods for peptide backbone generation and can be combined with such methods to rapidly find sequences predicted to fold correctly for a range of topologies.

### *De novo* Hallucination for Cyclic Peptides

We next developed a hallucination approach for cyclic peptides that samples the sequence and structure simultaneously and applied it to enumerate macrocyclic peptides beyond the helix-containing 13-mer scaffolds generated using our redesign approach. We implemented a three-step hallucination pipeline for generating sequences that are predicted to fold into structured cyclic peptides with high-confidence (see Methods). The approach is guided by losses that try to improve the prediction confidence metrics, pLDDT and predicted alignment error (PAE), and the number of intramolecular contacts.

We started with macrocycles composed of 7–10 amino acids and enumerated 48,000 hallucinated models for each size. We clustered the resulting structures from these large sampling runs using torsion bin-based clustering described earlier, and identified 9941, 13405, 19705, and 22206 unique structural clusters for 7-mers, 8-mers, 9-mers, and 10-mer cyclic peptides, respectively (Figure 3A). Out of all the unique structural clusters, 182, 297, 457, and 1282 clusters for 7-mers, 8-mers, 9-mers and 10-mers, respectively, had at least one member predicted to fold into the designed structure with high confidence (pLDDT > 0.9) (Figure 3A). Given our results on native structure prediction and redesign, we expect peptides that pass this strict confidence metric cutoff of 0.9 to fold correctly into the designed structure. We selected these sequences for further *in silico* validation of folding propensity by an orthogonal approach relying on Rosetta cyclic peptide structure prediction methods (*18, 20*) (see Methods for details). To evaluate the folding propensity of these sequences we calculated the P_near_ values as described in our previous work (*18, 20*). P_near_ values range from 0 to 1, and a value of 1 indicates that the designed structure is the single lowest energy conformation for that sequence (*20*). Many of these hallucinated sequences demonstrated promising folding propensity in these calculations, with 114 7-mers, 186 8-mers, 139 9-mers, and 76 10-mer sequences showing Rosetta P_near_ values greater than 0.6. We selected one hallucinated design model per size between 7–10 amino acids with AlphaFold pLDDT > 0.9 and Rosetta P_near_ > 0.9 for further experimental validation and structural characterization. All four selected designs lack regular secondary structures but are stabilized by extensive intramolecular backbone-to-backbone and backbone-to-sidechain hydrogen bonding. The design models for RH7.1, RH8.1, RH9.1, and RH10.1 feature 3, 5, 5, and 6 intramolecular hydrogen bonds, respectively (Figure 3, second column). The overall shape of the selected models is also guided by combinations of canonical α, β, and γ turns. Design RH7.1 is composed of a type I β turn and an overlapping γ and α turn, with all turns nucleated by proline residues. Design RH8.1 includes two type I β turns stabilized by a sidechain-to-backbone hydrogen bond from the aspartate residue at *i* position to NH of *i* + 2. Design RH9.1 also contains two type I β turns separated by a pair of long range hydrogen bonds between methionine-4 and leucine-9. Design RH9.1 is notable in its hydrophobicity, with the only polar residue in the design being a single glutamate. The sequence for RH10.1 is also significantly hydrophobic with multiple exposed nonpolar sidechains and hydrophobic packing between the tryptophan and a leucine stabilizing a region with few intramolecular hydrogen bond interactions. We also observed multiple glycines and prolines in the selected design models, with prolines providing the conformational constraints and glycines accessing the X and Y bins (phi angle > 0 degrees) of the Ramachandran plot.

**Figure 3:**
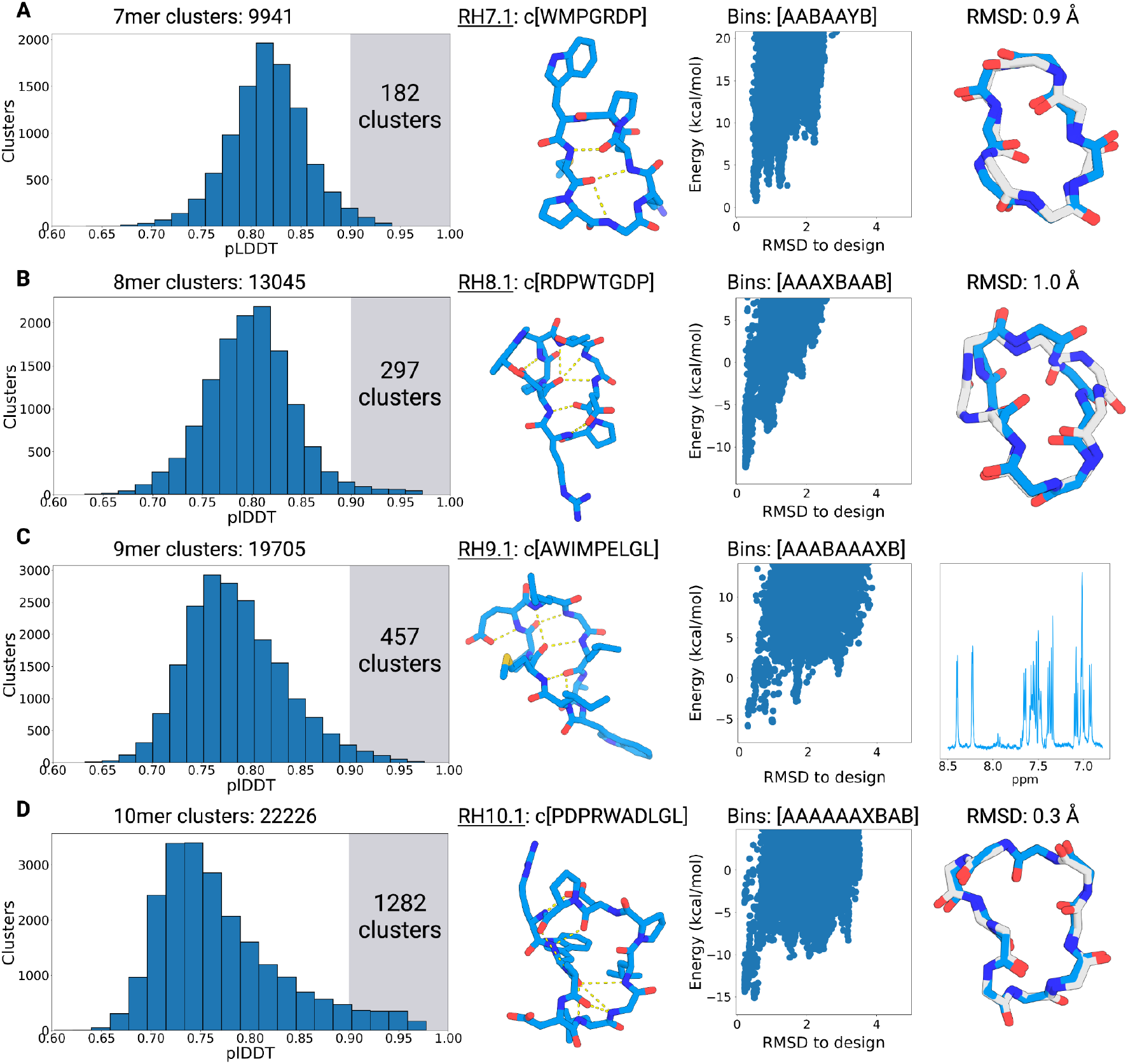
Hallucinating 7–10 residue cyclic peptides using AfCycDesign. Distribution of structure prediction confidence metrics (pLDDT) and validation of selected candidates from the large-scale sampling of (A) 7-mers, (B) 8-mers, (C) 9-mers, and (D) 10-mers. For each row, the first column describes the distribution of pLDDT scores for all unique structural clusters identified from 48,000 hallucinated peptides for sizes 7-10 residues. The total number of unique clusters for each size is described in the plot title. The highlighted area in each plot shows the number of clusters with pLDDT scores > 0.9. The <u>second column</u> shows the hallucinated structures and sequences for models selected for structural characterization. Hydrogen bonds are denoted by the dashed yellow lines. The third column shows the Rosetta calculated energy landscapes for the selected hallucinated models. Each scatter dot (blue) denotes a different conformation for the same designed sequence. The torsion bin string for the selected design model is shown on top of the plot. The <u>fourth column</u> shows the alignment between the hallucinated model (blue) and the X-ray crystal structure (gray). 1D NMR spectrum is shown for RH9.1.

We attempted to crystallize all four selected peptides, and obtained high-resolution x-ray crystal structures for the 7-mer, 8-mer, and 10-mer peptides (Figure 3, fourth column). The X-ray structures for the RH7.1 matched very closely with the hallucinated model, with a Cα RMSD of 0.9 Å between the two structures. There were small differences between the design model and X-ray crystal structure, with the X-ray crystal structure featuring an additional hydrogen bond from an aspartate sidechain stabilizing the type I β turn that was not designed. Comparatively, the RH8.1 structure deviated more from its hallucinated model with Cα RMSD of 1.0 Å and torsions around Proline-3 and Glycine-6 differing between the designed model and X-ray crystal structures. The differences are most striking at the singular glycine position in the sequence, where the ɸ torsion is flipped. RH10.1 structure is almost identical to the hallucinated model with Cα RMSD of 0.3 Å. The sidechain rotamers in the crystal structure also match the design model remarkably well, with the two leucines and the aspartate being identical. The □1, □2 and □3 dihedral angles of the arginine rotamer also match well, only deviating to form a salt bridge with the aspartate. We were unable to get an X-ray structure for the RH9.1; however, the 1D NMR for that design shows sharp and dispersed peaks suggesting it is also folded (Figure 3C, fourth column). Further structural characterization is required to confirm if the folded state matches the design model.

Next, we focused on hallucinating larger macrocycles composed of 11–13 amino acids. In our previous attempts with Rosetta-based approaches, designing large structured macrocycles posed significant challenges and required additional disulfide crosslinks to stabilize them (*18*). We wondered whether AfCycDesign could hallucinate macrocycles in this size range without requiring additional crosslinks. For 11-mer, 12-mer, and 13-mer macrocycles, we identified 28457, 27715, and 27056 unique structural clusters, respectively (Figure 4, first column) from large-scale design calculations (see methods). We had a considerable number of clusters with members predicted to fold into the designed structures with high confidence: 1810, 2855, and 3798 clusters for 11-mer, 12-mer, and 13-mer, respectively, had members with AlphaFold pLDDT > 0.9. We selected one sequence with pLDDT > 0.9 and Rosetta P_near_ > 0.9 for each size for experimental validation. In contrast to the smaller 7–10 amino acid design models, we noticed short motifs of canonical secondary structures in the selected three designs from 11–13 amino acids (Figure 4, second column). Design RH11.1 contains an eight residue α-helical motif and RH12.1 and RH12.1 feature short extended β-sheets. Notably, with sequence lengths of 12 and 13 amino acids, both RH12.1 and RH13.1 have atypical sizes for cyclic beta strands as they are more favored in sizes 6, 10, and 14 residues (*28*). The larger structures in these selected peptides also include extensive intramolecular hydrogen bonds, with 9, 7, and 9 hydrogen bonds in the 11-mer, 12-mer, and 13-mer design models, respectively. The short helical motif in RH11.1 is cyclized via a four-residue extended loop and features an N-terminal helix-capping motif mediated by a threonine residue (Figure 4A, second column). RH12.1 is a short β sheet with a canonical type II’ β-turn connecting the strands on one end, and an α turn on the other end. In RH13.1, the register of the strand pairing is shifted by a cross-strand hydrogen bond from an aspartic acid sidechain to a backbone amide nitrogen, creating a twisted β-sheet cyclized at two ends by a type I β turn and an α turn, and further stabilized by hydrophobic interactions between the non-polar sidechains (Figure 4, second column).

**Figure 4:**
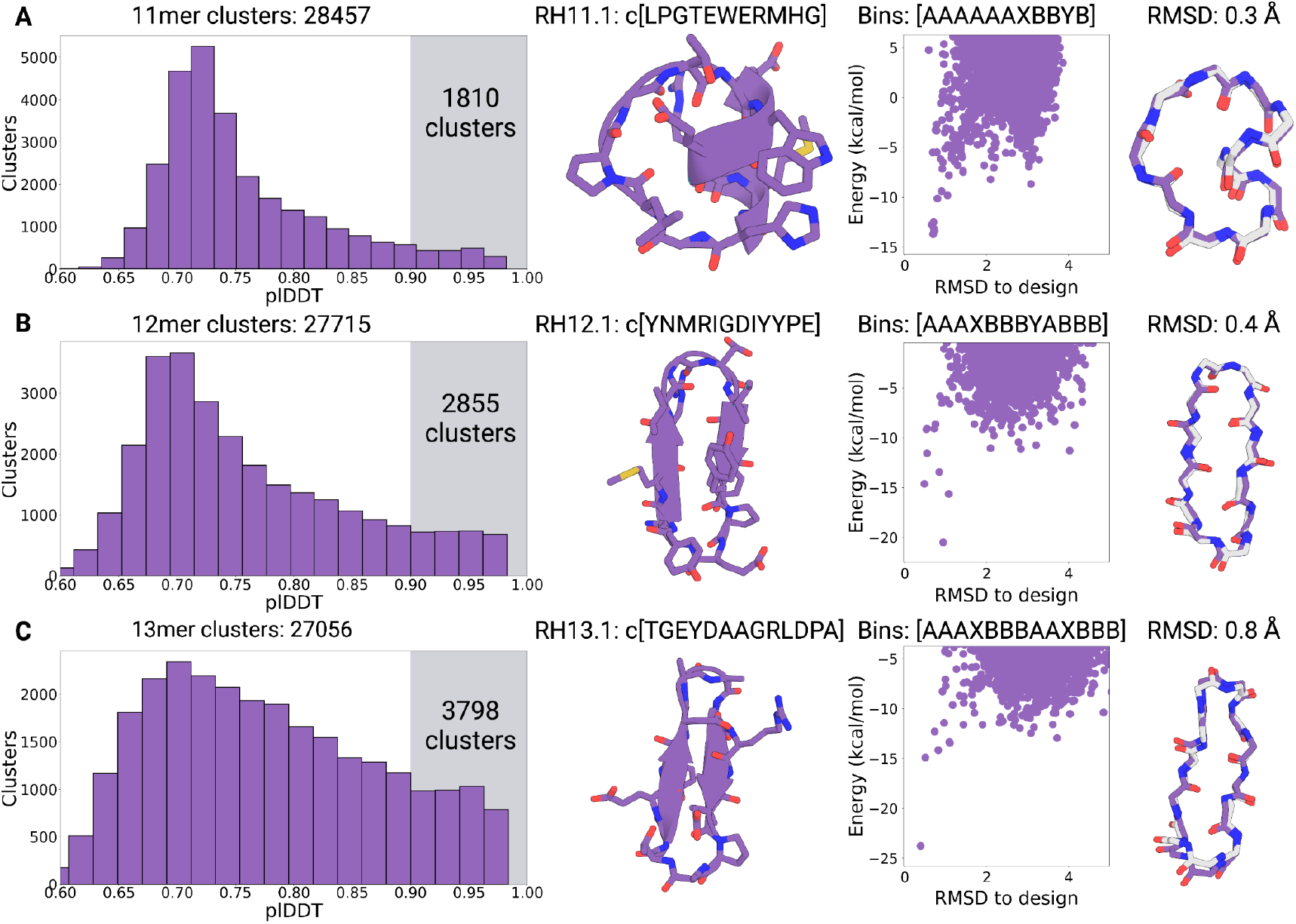
Hallucinating 11–13 residue cyclic peptides using AfCycDesign. Distribution of structure prediction confidence metrics (pLDDTs) and validation of selected candidates from the large-scale sampling of (A) 11-mers, (B) 12-mers, and (C) 13-mers. For each row, the first column describes the distribution of pLDDT scores for all unique structural clusters identified from 48,000 hallucinated peptides for sizes 11-13 residues. The total number of unique clusters for each size is described in the plot title. The highlighted area in each plot shows the number of clusters with pLDDT scores > 0.9. The second column shows the hallucinated structures and sequences for models selected for structural characterization. The third column shows the Rosetta calculated energy landscapes for the selected hallucinated models. Each scatter dot denotes a different conformation for the same designed sequence. The torsion bin string for the selected design model is shown on top of the plot. The fourth column shows the alignment between the hallucinated model (purple) and X-ray crystal structure (gray)

We synthesized RH11.1, RH12.1, RH13.1, and their mirror images, using solid-phase chemical peptide synthesis, and determined the structures for all three peptides using racemic X-ray crystallography. High-resolution crystal structures for all three peptides matched very closely with their hallucinated models, with Cα RMSDs of 0.3, 0.4, and 0.8 for the RH11.1, RH12.1, and RH13.1, respectively (Figure 4, fourth column). The turn types and hydrogen bonding patterns in the X-ray crystal structures also match closely with the design models for all three peptides. The greater RMSD observed in RH13.1 is due to small torsional deviations propagating along the peptide chain as opposed to changes in turn types or flipped amides previously seen for RH7.1 and RH8.1. Most of the critical sidechain interactions in the designs are also observed in the X-ray structures; the two most obvious deviations being the tryptophan in RH11.1 flipping 180°, but still ring stacking with histidine as seen in the design, and a tyrosine in RH12.1 that instead of making interactions with the backbone has rotated up to form a cation-π interaction with an arginine. Taken together, these data highlight the excellent accuracy of AfCycDesign for *de novo* hallucination of cyclic peptides, including larger macrocycles between 11–13 amino acids without requiring additional disulfide bonds to stabilize such structures as was previously proposed. More broadly, the hallucination approach and extensive structural sampling described here provide several new scaffolds for incorporating functions.

## CONCLUSION

We report an approach to incorporate cyclic relative positional encoding in the AlphaFold network and leverage it to develop computational methods for several key applications, including, structure prediction of cyclic peptide sequences, redesigning amino acids on natural and designed cyclic peptide backbones, and *de novo* hallucination of new cyclic peptides with diverse sequences, sizes, and topologies. Our tests with structure prediction of previously described cyclic peptides highlight the remarkable accuracy of AlphaFold with cyclic offsets; 50 of the 80 sequences were predicted correctly with RMSD to the native structure < 1.5 Å and pLDDT > 0.7. Among the 49 designs that were predicted with high confidence (pLDDT > 0.85), 73% match the NMR-detemined structures with RMSD < 1.5 Å. The accurate structure predictions from our approach should enable rapid and reliable structural insights for naturally-occurring cyclic peptides, and enable better filtering for computationally-designed peptides expected to fold correctly into the designed structures.

We also describe AfCycDesign, a computational method to redesign sequences of cyclic peptide backbones, with AfCycDesign sequences showing better pLDDTs and folding propensities than sequences generated by the previously described Rosetta sequence design approach for the same peptide backbones. Comparing the AfCycDesign sequences and Rosetta-designed sequences from a large-scale redesign of 13-mer cyclic peptides highlights some key differences, including increased usage of hydrophobic and conformationally-restricted amino acids by AfCycDesign. We further extended our AfCycDesign approach for hallucinating sequence and structures for cyclic peptides simultaneously and applied it to enumerate hundreds of thousands of unique structural clusters for peptides composed of 7–13 amino acids, including 10,681 unique clusters that are predicted to fold into the designed structures with very high confidence (pLDDT > 0.9). X-ray crystal structures for the redesigned and hallucinated cyclic peptides show notable accuracy of our methods: All seven x-ray crystal structures (one redesigned and six hallucinated) are remarkably close to their design models with RMSDs less than 1.0 Å. Notably, the hallucination approach allowed us to successfully design larger cyclic peptides between 11–13 amino acids that had proven difficult to design without additional crosslinks in previous attempts with state-of-the-art approaches (*18*). Further, we previously noted the importance of L- and D-amino acid patterning in generating structured cyclic peptides (*18*); however, the hallucinated peptides described here defy those guidelines and are well-folded despite being composed of L-amino acids only. We acknowledge the protease and metabolic stability benefits provided by D-amino acids and other non-canonical amino acids, and believe this work provides the basis for developing deep learning networks in future that can incorporate a broader chemical diversity during design.

Hallucinated peptides predicted to fold into their designed structures with high confidence (pLDDTs > 0.9), and their mirror images, should serve as excellent scaffolds for incorporating functions, such as target binding and membrane traversal. Binding interactions could be incorporated by grafting motifs from protein-protein interactions or designed *de novo*. A focus of our current and future efforts is to extend the computational approach for hallucinating cyclic peptide binders against therapeutic targets. Deep learning methods have led to large advances in therapeutic protein design over the last five years. With the computational approach presented here, similar advances can be extended to the custom design of structured cyclic peptides with high therapeutic significance.

## METHODS

### Cyclic Peptide Design with AfCycDesign

The protocol, implemented within the ColabDesign v1.1.0 framework, is used to generate a protein sequence that either folds into a desired target backbone structure or to hallucinate a new protein. Since a discrete sequence is not differentiable, we use the 3-stage design protocol that starts from a continuous representation and eventually ends with a one-hot encoded sequence. The input sequence is represented as a peptide length × 20 matrix. The sequence-matrix starts as an unconstrained and continuous set of logits and gradually becomes a normalized probability distribution using a formula that combines logits and softmax probabilities: ((1-p) * logits + p * softmax(logit/temperature)). In the first 300 iterations, the formula emphasizes logits more, but the emphasis shifts towards softmax probabilities over the iterations until a softmax distribution is achieved in stage 2. The temperature is then reduced in the next 200 iterations, approaching a one-hot encoded sequence. More specifically, p is linearly scaled from 0 to 1 in stage 1, and temperature is reduced from 1.0 to 0.01. In the third and final stage, the one-hot encoded sequence is directly optimized for 10 steps using a straight-through estimator. Throughout the optimization process, dropouts are enabled, and the model parameters are randomly selected from three models to escape local minima. At the third stage, the dropouts are disabled, and the sequence with the best loss is chosen as the final design.

For fixed backbone (fixbb) design, the categorical-cross-entropy (CCE) loss between the desired and the predicted distogram is used. For hallucination design, a combo of three losses is used. This includes 1-pLDDT + PAE/31 + con/2. pLDDT and PAE are average confidence metrics returned by AlphaFold. The [con]tact loss was designed to maximize the number of interacting residues, designed to promote a compact structure. For peptide design, the default contact loss was modified using the following settings: (binary=True, cutoff=21.6875, num=length, seqsep=0). To promote structural diversity, we initialize the sequence with a random Gumbel distribution. The first 50 steps of optimization are primed with softmax activation and temperature of 1.0. Standard offset matrix is used for the relative positional embedding. The cyclic offset is then enabled, sequence is initialized with the softmax(logits), and the 3-stage protocol, with schedule of (stage1=50, stage2=50, stage3=10), is run to get the final one-hot sequence.

### Energy landscape calculation and analysis

Energy landscape calculations for cyclic peptides were done as previously described using Rosetta cycpep_predict application (*18–20*). Large-scale conformational sampling during these calculations was conducted using the BOINC Rosetta@Home platform. Energy for each sampled conformation was calculated using the Rosetta REF2015 energy function(*29, 30*). The folding propensity was evaluated based on the energy gap between the design conformation and alternative conformations, and by calculating P_near_, a Rosetta metric that looks at the quality of energy ‘funnel’. P_near_ value of 1 denotes energy landscapes with a funnel that converges to the designed model as its single low-energy minima; 0 denotes energy landscapes with one or more alternative conformations as energy minima that are different from the designed conformation. See ref (*20*) for details of the P_near_ metric.

### Peptide Synthesis and Purification

Macrocyclic peptides were purchased from Wuxi AppTec at greater than 90% purity or synthesized in-house by manual Fmoc-based solid-phase peptide synthesis at 0.2 mmol scale. 300 mg of 2-Cl-Trt resin purchased from Anaspec was transferred to a 10 ml Torviq disposable reaction vessel and swelled for 1 hour in dichloromethane (DCM). Linear peptide synthesis was initiated on glycine residues found in the sequences, 0.2 mmol of Fmoc-Glycine-OH in 5 ml of DCM with 300 μl of 2,4,6-collidine was incubated overnight with the resin. The resin was drained and washed 3X with DCM followed by capping with a 17:2:1 mixture of DCM:methanol:DIEA (N,N-Diisopropylethylamine) for 1 hour. After capping, the resin was washed 3X with DCM and 3X with N,N-Dimethylformamide (DMF) and subjected to repeated 20 minute deprotections with 20% piperidine in DMF followed by 3X washes with DMF and 20 minute couplings with 5 eq of Fmoc protected amino acid mixed with 5 eq of PyAOP ((7-Azabenzotriazol-1-yloxy)trispyrrolidinophosphonium hexafluorophosphate) and 10 eq of DIEA dissolved in DMF. After removal of the terminal Fmoc group, the protected peptide was released from the resin by repeated washes of 2% TFA (trifluoroacetic acid) in DCM that were deposited in a round bottom flask containing a 50:50 mixture of water and acetonitrile. DCM was removed from the mixture by rotary evaporation and the resulting protected peptide, now in water and acetonitrile, was lyophilized to dryness. The dry peptide was dissolved in 50 ml of DCM with 2 eq of PyAOP in a round bottom flask with a stir bar and left to stir for 10 minutes. 3 eq of DIEA was added dropwise and the solution left overnight. DCM was removed by rotary evaporation leaving an oil-like solution in the round bottom flask. To remove the protecting groups, 20 ml of TFA:water:triisopropylsilane:3,6-Dioxa-1,8-octane-dithiol (92.5:2.5:2.5:2.5) was added and the mixture left stirring for 3 hours. The deprotected peptide was concentrated by rotary evaporation, precipitated in cold diethyl ether, and dried under air stream. Peptides were purified on an Agilent Infinity 1220 HPLC using 1% per minute gradient on Agilent ZORBAX SB-C18, 80Å, 5 mm, 9.4 × 250 mm column with a gradient of solvent A: 0.1% TFA in water, and solvent B: 0.1% TFA in acetonitrile. Mass spectrometry was used to confirm the synthesis of the correct product; purified peptides were direct-injected on a Bruker Esquire Ion Trap Mass Spectrometer.

### X-ray Crystallography

Crystals diffraction data were collected from a single crystal at synchrotron (on APS 24ID-C) and at 100 K. Unit cell refinement, and data reduction were performed using XDS and CCP4 suites (*31, 32*). The structure was identified by direct methods using SHELXT (*33*). RH7.1 was refined by full-matrix least-squares on F^2^ with anisotropic displacement parameters for the non-H atoms using SHELXL-2018/3 (*33*). Structure analysis was aided by using Coot/Shelxle (*34, 35*). The hydrogen atoms on heavy atoms were calculated in ideal positions with isotropic displacement parameters set to 1.2 × U_eq_ of the attached atoms. All structures deposited in CDS/CCDC (The Cambridge Structural Database/Cambridge Crystallographic Data Centre).

### NMR Spectroscopy

Peptide sample for NMR data collection was prepared by dissolving 5-10mg of peptide in 600 microliter solution of 20:80 CD_3_CN:H_2_O. The NMR data was collected using Bruker Avance II 500 MHz spectrometers equipped with TCI cryoprobes. The data was collected at 293 K and referenced to internal TMS. The spectrum was processed using MestReNova (*36*). Chemical shifts are represented in parts per million (ppm).

## ACKNOWLEDGEMENTS

We thank David Baker, Lance Stewart, Justas Dauparas, Lauren Carter, Luki Goldschmidt, Preetham Venkatesh, Patrick Salveson, Meerit Said, Ian Haydon, Paul M. Levine, Xinting Li, Mila Lamb, Martin Sadilek, Rajan Paranji, Theresa Ramelot, and members of the Bhardwaj lab and the Institute for Protein Design for helpful discussions. We thank the volunteer contributors of the BOINC Rosetta@Home project for donating compute cycles for this project. We also thank the IPD core labs, the University of Washington (UW)’s Chemistry NMR facility, and the UW Chemistry mass spectrometry facility for providing instrumentation support and expertise. G.B. is supported by funds from DARPA Harnessing Enzymatic Activity for Lifesaving Remedies (HEALR) program (HR001120S0052 contract HR0011-21-2-0012), the Defense Threat Reduction Agency (DTRA) (HDTRA1-19-1-0003), Howard Hughes Medical Institute (HHMI) Emerging Pathogens Initiative, Bill and Melinda Gates Foundation (OPP1156262 Macrocycles), and start-up funds from UW Medicinal Chemistry and the UW Institute for Protein Design. S.O. and S.K. were supported by NIH DP5OD026389, NSF MCB2032259, and Amgen. Crystallographic data was collected at the Advanced Photon Source (APS) Northeastern Collaborative Access Team beamlines, which are funded by the National Institute of General Medical Sciences from the National Institutes of Health (P30 GM124165). This research used resources from the Advanced Photon Source, a U.S. Department of Energy (DOE) Office of Science User Facility operated for the DOE Office of Science by Argonne National Laboratory under Contract No. DE-AC02-06CH11357. All plots were generated using matplotlib (*37*). Peptide structures were rendered using PyMOL, and figures were created using BioRender.

## CODE AVAILABILITY

Example scripts for structure prediction, sequence design, and hallucination are available at https://github.com/sokrypton/ColabDesign/blob/main/af/examples/af_cyc_design.ipynb Rosetta software suite can be downloaded from https://www.rosettacommons.org/

---

**Figure S1:**
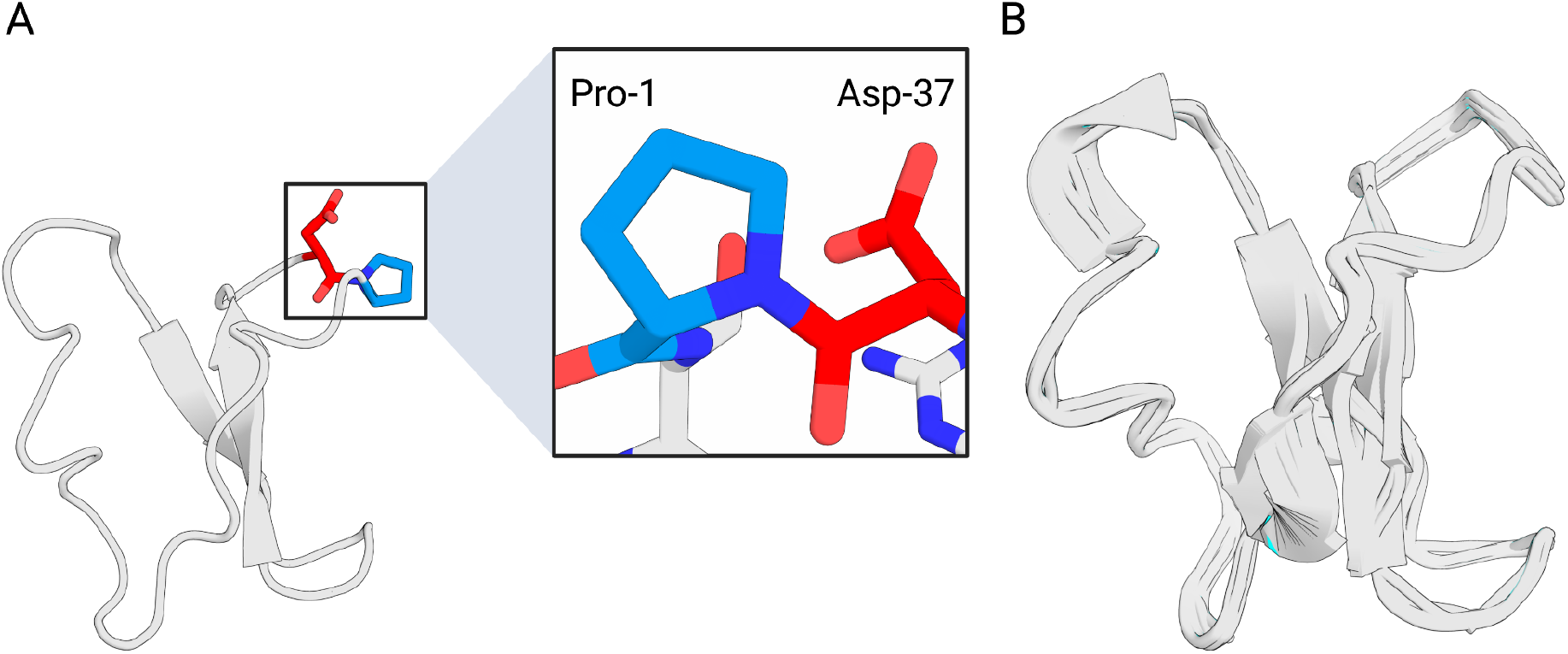
Cyclic offset to relative positional encoding in AfCycDesign enforces N-to-C terminal cyclization and is invariant to circular permutations of the sequence. (A) Peptide structures predicted with AfCycDesign show correct bond connectivity and geometry at the termini. (B) Cartoon overlay of 37 predicted models for each circularly permuted sequence of a native peptide with cyclic offset applied in AfCycDesign.

**Figure S2:**
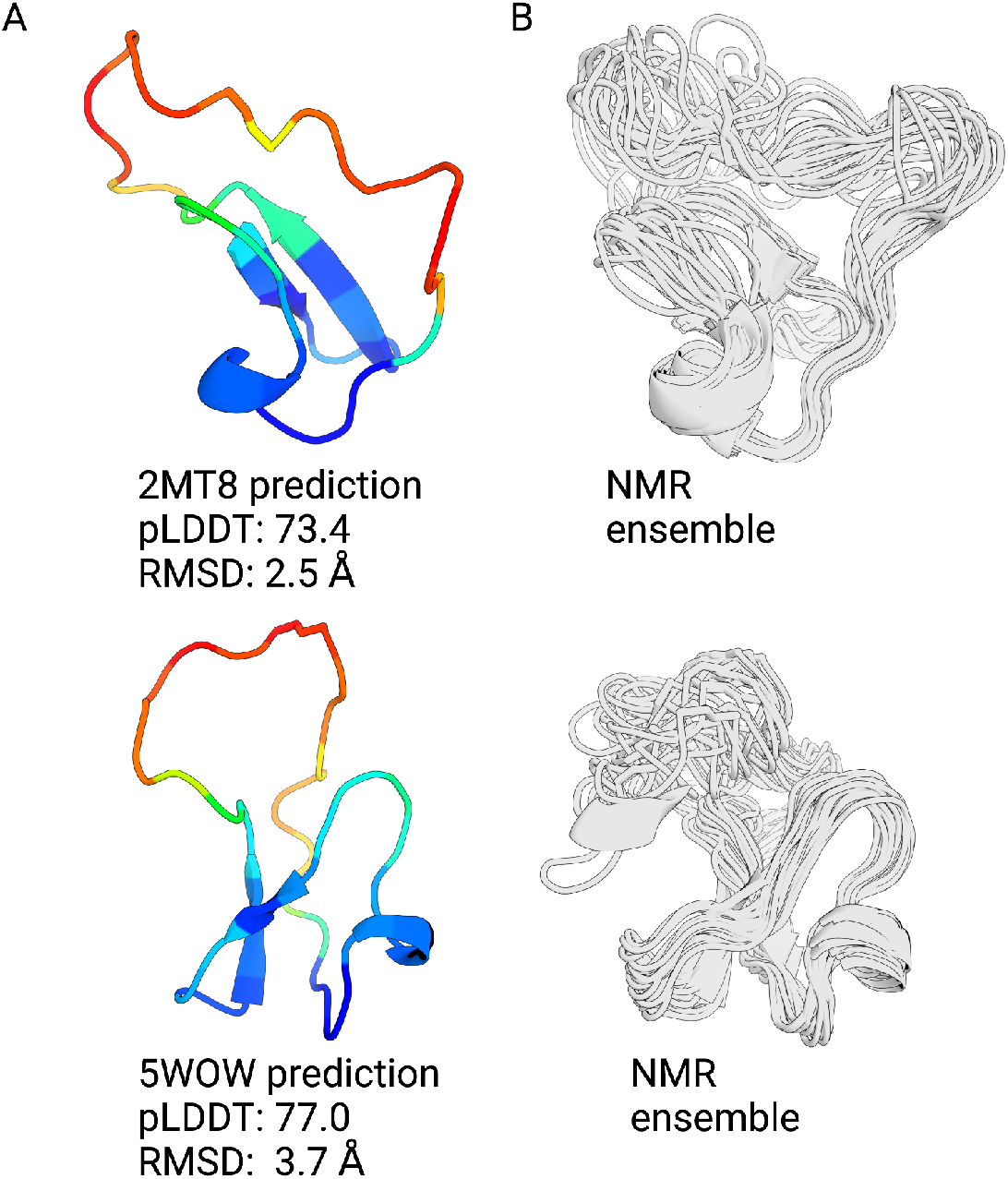
Regions predicted with low confidence match the flexible regions in the NMR structures. (A) Representative peptides with pLDDT lower than 0.85. Cartoon representations are colored from regions of high confidence (blue) to low confidence (red) based on per residue pLDDT. RMSD is calculated against all ensemble members, and the lowest backbone heavy atom RMSD is reported. (B) NMR ensemble of the peptide aligned to the predicted model.

**Figure S3:**
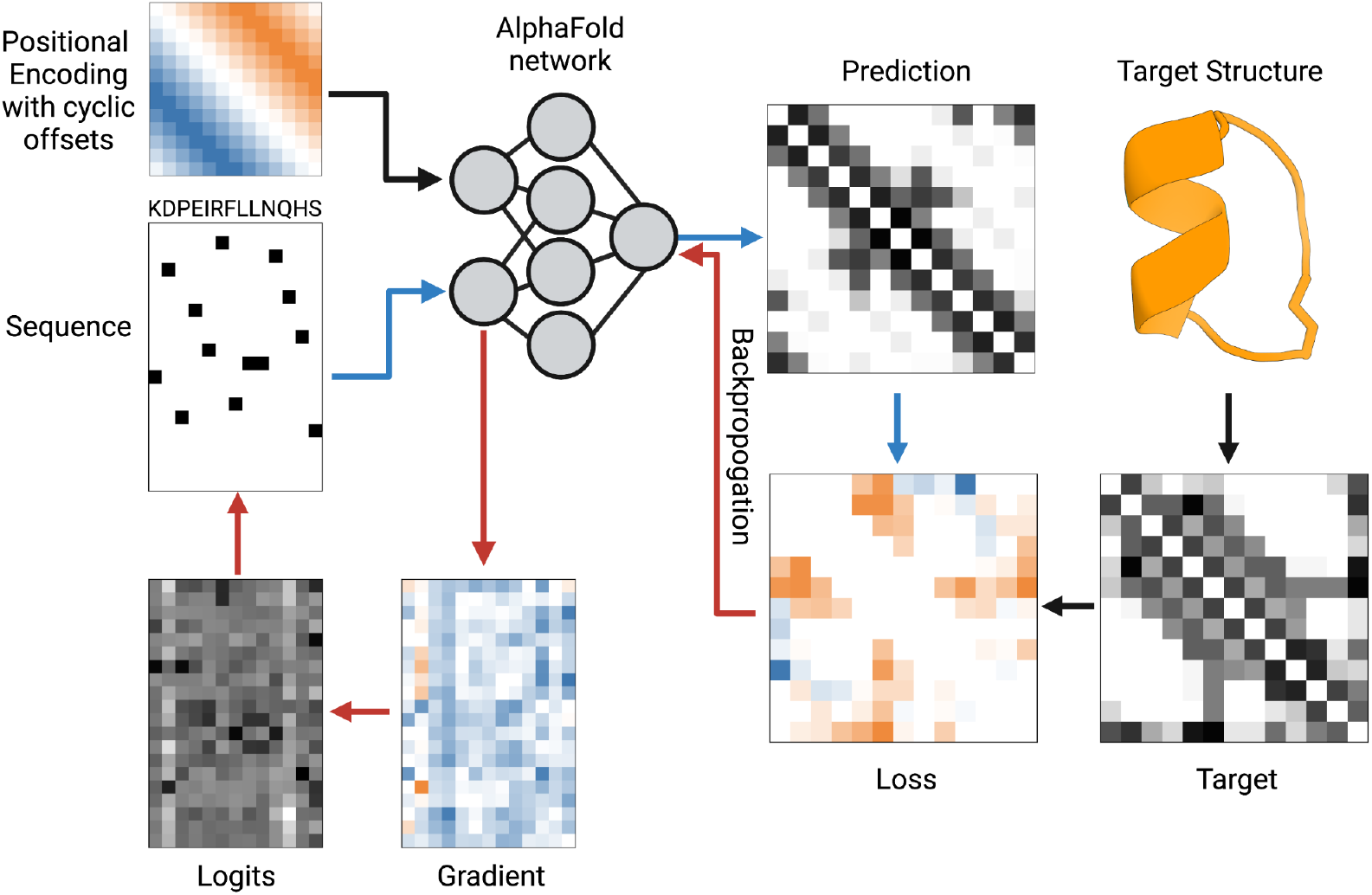
Overview of the sequence design approach in AfCycDesign. Given the input sequence and relative positional encoding with cyclic offset, AlphaFold is used to predict the distogram (distribution of distances between every pair of positions). The loss is defined as the categorical-cross entropy (CCE) between the output distogram and the desired distogram extracted from the desired structure. At each iteration, the gradient is computed to minimize the CCE. The gradient is then applied to a proxy variable we term “logits”. The one-hot encoding of the argmax of the logits is then used as the input sequence for the next iteration.

